# Comparative Analysis of Cameras and Software Tools for Skeleton Tracking

**DOI:** 10.1101/2023.08.10.552434

**Authors:** Naama Aharony, Amir Meshurer, Maya Krakovski, Yisrael Parmet, Itshak Melzer, Yael Edan

## Abstract

**Background:** Skeleton tracking is a valuable tool for monitoring user performance and has been performed with different sensors and software tools. It enables to quantify 3D motion accurately. We present a comparative analysis of cameras and software tools utilized in skeleton tracking. We describe an experimental design and analysis methods with four steps. Specifically, we conducted an analysis comparing two ZED-2i cameras with a RealSense camera using commercial skeleton tracking algorithms.

**Methods:** The comparative analysis methodology describes the experimental design and analysis methods for comparing kinematic data from both upper and lower extremities. An experiment to evaluate the performance of three different cameras (RealSense, ZED2mm, and ZED4mm) and three skeleton tracking algorithms (PyZED, Nuitrack, and MediaPipe) was performed with 16 participants. The participants were required to walk along a 6-meter-long path while moving their hands up and down horizontally. Ground truth was measured by a Vicon 3D motion analysis system. Two groups of features corresponding to the upper and lower extremities were derived from the raw data and compared using two methods. In the first method, the RMSE values were compared to the data from the Vicon system. The RMSE results were also combined using two grading techniques with different feature importance weights to determine the camera with the lowest RMSE. In the second method, a linear mix model with a Wald statistical test was applied.

**Results:** The ZED-2i cameras outperformed the RealSense camera across upper- and lower-extremity features and different weights. The ZED2mm camera showed better average median grading, lower standard deviation, and superior performance in the model analysis.

**Conclusion:** The comparative analysis methodology provides a systematic method to compare software tools and cameras for skeleton tracking. The ranking enables to prioritize different features. Skeleton tracking with the ZED cameras yielded better performances for both upper and lower extremities features.

## 1 Background

Motion capture (MoCap) is the process of tracking and recording the movement of an object, enabling the user’s performance to be monitored [1]. This technology is integrated into many applications, such as rehabilitation [2–4] and physical training [5–7] and even in movies, animations, video games, and among high level athletes [8]. Skeleton tracking is a specific type of motion capture that uses sensors, frequently cameras, or depth cameras [9]. It involves tracking a human body’s or animal’s movement by identifying and tracking the locations of key joints or points on the body, such as the head, shoulders, elbows, and knees [10–12]. Once those joints are identified, the software connects them to a humanoid skeleton and tracks their position rein all time.

One technology that offers full-body skeleton tracking for three-dimensional (3D) motion analysis in fine detail, the Vicon 3D motion analysis system, utilizes multiple cameras and tracking markers placed on the skin. The system is expensive and is limited to lab environments [10,15]. Furthermore, this technology requires extensive expertise to properly analyze and interpret the data [8].

The advent of video-based pose tracking provides an opportunity for inexpensive automated analysis of human activity using video and depth cameras [13]. This equipment provides affordable and more accessible solutions that are easy to use and do not require markers to determine anatomical landmarks or extensive technical expertise to operate and interpret without sacrificing accuracy [14]. Low-cost 3D cameras such as Kinect, RealSense, and ZED-2i enabling applications to be developed that rely less on a technological background for real-life environments [16]. Kinect and RealSense have been previously validated and show potential for use in skeleton tracking [5,17,18]. However, Microsoft has stopped production of the Kinect camera [19], and Intel has announced that the RealSense product line will not continue [20].

Comparison between cameras and software tools for skeleton tracking was conducted in several previous studies [21], including pose tracking performance for different Kinect cameras [22,23], comparing stride-to-stride gait variability using vision and Kinect sensing [24–26], and comparing a new 3D marker-less motion capture technique using OpenPose with multiple synchronized video cameras to an optical marker-based motion capture [27]. The accuracy of depth cameras in body landmark location estimation was only measured for static uprights in [28].

This research focused on presenting a comparative analysis methodology of different cameras with skeleton tracking software for both groups of upper and lower extremities features. These features can be used in different applications, e.g., for fall detection, exercise feedback, and medical screening tests such as balance and gait assessment. Upper extremities applications include recognizing arm and hand motions [29] during rehabilitation exercises [2–4,30,31] and Arthritis management [32]. Lower extremities examples include gait event detection during curved walking and turning in older adults and Parkinson’s Disease patients [33,34], and estimation of ground reaction forces during balance exergaming [14]. The presented comparative analysis methodology provides a systematic way to compare software tools and cameras for skeleton tracking, enabling to prioritize different features depending on the specific research objectives.

## 2 Methods

We propose the following four steps for comparative analysis of different cameras and skeleton tracking software (Fig 1). The demonstration of these methods in a specific comparative analysis is described in the following section (section 3).

**Fig 1.**
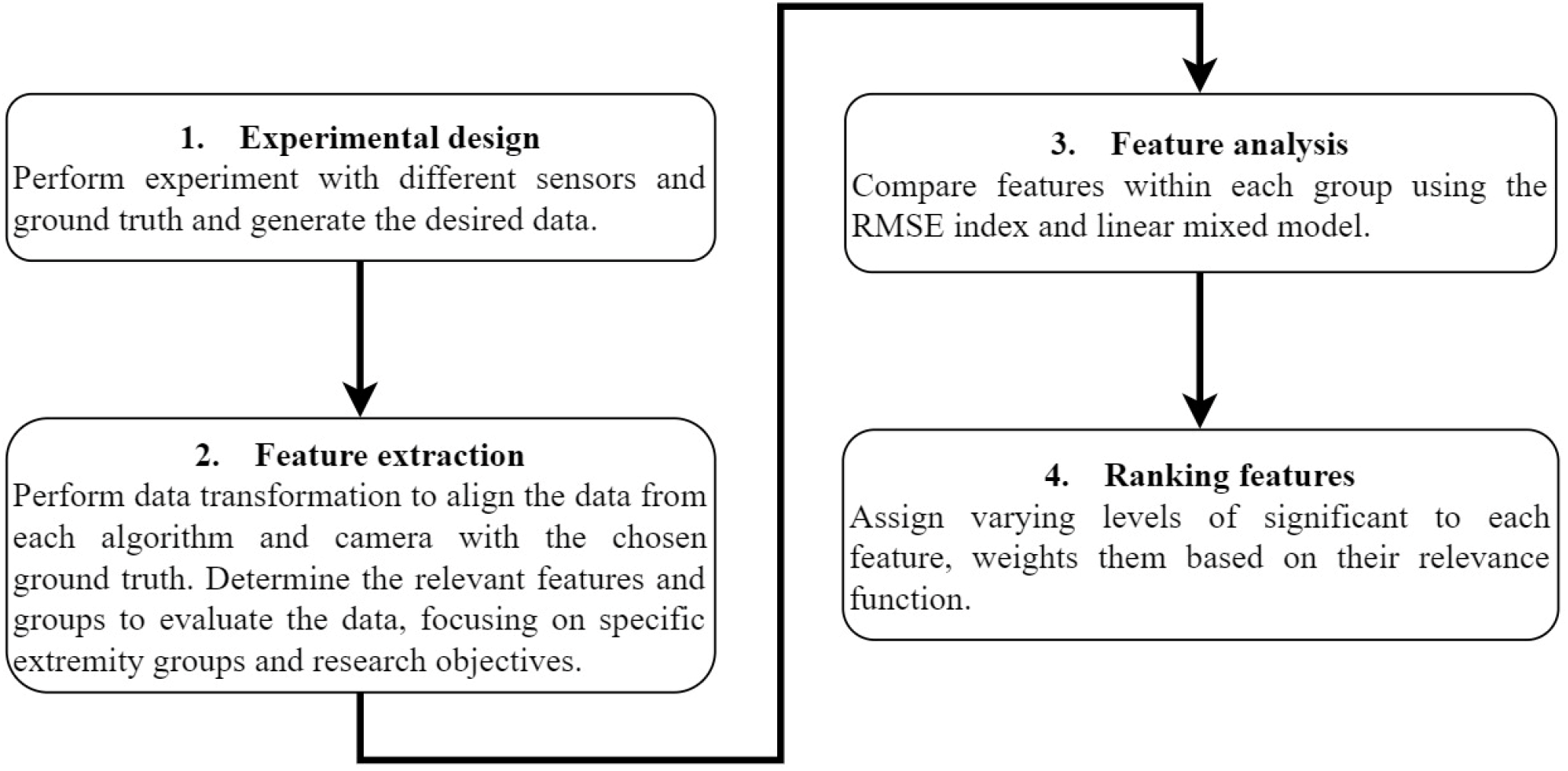
Comparative analysis flowchart.

### 2.1 Experimental design

The experimental design should include measurements of human motions with different sensors and a ground truth sensor. Cameras should be placed in the same position in all experiments. Since some cameras/algorithms require dedicated capture of the motion, simultaneous measurement of all systems in real-time is only sometimes possible due to hardware limitations. Although there are methods for accurate alignment without requiring the cameras to be synchronized and without aligning captured frames in time (e.g., [35] these usually require measuring the intrinsic and extrinsic camera parameters and calibration procedures which are complicated and have inherent errors [36–38]. Additionally, in dynamic motions in natural settings, there is inherent variability between participants and trials [21].

Since the aim of this research is comparative analyses and not the actual accuracy of these sensors, the experimental design includes independent analysis of each specific camera and algorithm. In each trial, a different camera with the proper algorithm is activated and compared to the ground truth measurement. To create a variety of conditions, multiple participants should conduct repeated trials. The sequencing of the trials between cameras and the algorithms must be randomized to ensure unbiased results.

### 2.2 Feature extraction

Due to the diverse hardware used for data collection in each camera and algorithm, direct comparison of raw data is impractical as continuous distributions (e.g., the time the data is sampled) rarely have identical values. To enable performance comparison, we must extract time-independent features from the raw data.

Data transformation is necessary to align the camera and algorithm data with the ground truth, so that each joint of each algorithm can be compared with the joint of the ground truth measurements.

Then, the relevant features should be calculated from the data. Every feature should be calculated for each specific trial and specific camera with the proper algorithm. These features should be assigned to groups, focusing on particular extremity groups according to specific research objectives.

### 2.3 Feature Analysis

Every feature for each specific trial and camera with the proper algorithm should be compared to the ground truth by the root of the mean square error (RMSE) index. Then, features within each group should be compared using the RMSE index and linear mixed model for compression analysis, as detailed below.

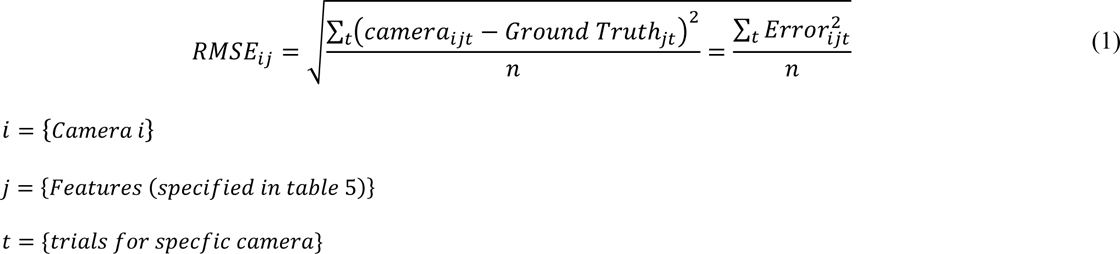

Additionally, a linear mixed model should be constructed for each individual feature to provide a better understanding of the results and examine the coefficients and determine whether the distributions are equal. This approach provides a tool to compare the ground truth features results to those obtained from the examined depth cameras with the different algorithms. Linear mixed models are a powerful tool for analyzing complex datasets and can be particularly useful in examining the coefficients and distributions of feature data. By considering both repeatable and non-repeatable covariates, these models can account for a wide range of factors that might affect the outcomes being studied [39,40].

For each feature and each camera with the proper algorithm, the model should be executed for both fixed and random effects. Fixed effects are those that remain constant throughout the analysis (repeatable covariate), while random effects are those that vary depending on the specific observation being studied, the participants’ effect in all trials - 𝑢_𝑠𝑢𝑏𝑗𝑒𝑐𝑡_ in equation 2 (non-repeatable covariate).

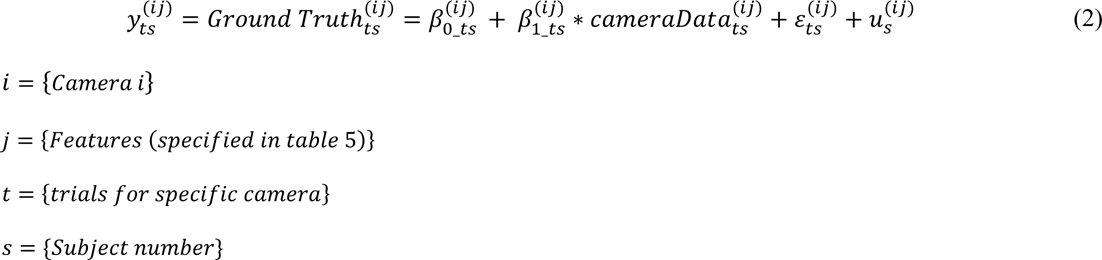

In Equation 2, 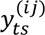 is the Ground Truth 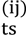 that was compared to the proper cameraData_ijt_ to check one on one relationship. 𝛽_0_ts_ represent the intercept, 𝛽_1_ts_ represent the regression coefficient (slope), 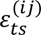 represent the noise of the data and 𝑢_𝑠_ represent the participant’s effect on the model (random effect).

To evaluate the coefficients obtained from the models in Equation 2, the Wald statistical test [41] was used. The test compares a model coefficient to a null hypothesis. By analyzing these coefficients, we can gain insights into how the features contribute to the outcome being studied and identified which factors were the most important. This test provides a measure of statistical significance that can help to determine whether the tested combination of depth camera and algorithms produced good results for a particular feature. Since the aim is to determine if the distributions are equal (One to one relationship), the null hypothesis is 𝜷_𝟏_ equals to one and 𝜷_𝟎_ represents a constant bias.

In a perfect world, the slope, 𝛽_1_, should be 1, and the intercept, 𝛽_0_, should be 0. If the slope is equal to 1, but the intercept is different from 0, then the camera’s bias relative to the round truth is constant regardless of the actual size. If the slope is different from 1, the bias depends on the measured value of the camera, 𝛽_0_ + 𝑐𝑎𝑚𝑒𝑟𝑎𝐷𝑎𝑡𝑎(𝛽_1_ − 1). In some sense, 𝛽_0_and 𝛽_1_can be considered calibration constants of the camera measurements to the ground truth. Mixed-effects models come in two types of *R*^2^, marginal and conditional. The marginal represents the variance explained by the fixed effects, and the conditional is interpreted as a variance explained by the entire model, including both fixed and random effects. Since our goal is to evaluate the model’s ability on new observations of new subjects and not on existing subjects, the marginal *R*^2^ will be of interest to us.

To summarize, the analysis of the linear mix model was conducted in two stages, both in Python and R Programming in RStudio (the link to the code is given in Appendix A1). The first stage was to construct the model and its *R*^2^s, and the second step was to perform the Wald test on 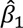, were the null hypothesis states that it is equal compared to one.

### 2.4 Ranking features

In order to compare different features with different scales the following should be attended:

- Significant weights - Assign varying levels of significant to each feature, weights them based on their relevance to the research/objective function.
- Grading techniques - Utilize different grading techniques to ensure equal scales when comparing features using the RMSE index.

## 3 Comparative analysis demonstration

To demonstrate the proposed comparative analysis, an experiment was conducted with three different cameras (ZED2mm, ZED4mm, and RealSense) and three algorithms (PyZED, Nuitrack, and MediaPipe) for skeleton tracking of upper and lower extremity features. This section describes the four steps of the comparative analysis based on our experiment.

### 3.1 Experimental design

The first step in the comparative analysis includes the technical specifications of the systems, followed by the experimental protocol, setup, and procedures.

#### 3.1.1 Technical background

<u>Hardware:</u> (Table 1)

- Intel’s RealSense D-435 camera [43];
- ZED-2I camera by StereoLabs, which has two kinds of lenses, 2mm and 4mm [44]; Ground truth was measured by a Vicon system (Vicon Motion Systems, Oxford, UK) [45–47].

**Table 1.** Technical specifications of the cameras compared and the ground truth Vicon system.

| Parameter | Intel RealSense D435 | ZED-2i | Vicon system |
| --- | --- | --- | --- |
| Technology | Active stereoscopy | Stereo vision and neural networks | Optical motion capture |
| Depth range (m) | 0.2m–10m | 0.2m–20m | 0.5m–12m |
| RGB resolution (pixel) | 1980 X 1080 | 3840 X 1080 | 8 Megapixel (8,000,000) |
| Depth resolution (pixel) | 1280 X 720 | 3840 X 1080 | 2432 X 2048 |
| Field-of-view depth<br>(FOV) | 85.2 X 58 | 110(H) X 70(V) X 120(D) | 100.6 X 81.1 |
| Frames per second (FPS) | 90 FPS | 30 FPS | 330 FPS |
| Weight (g) | 100 g | 166 g | 570 g |
| Cost (\$) | 179 \$ | 500 \$ | 30,000+ \$ |

<u>Software:</u> The RealSense (RS) camera was tested with two different algorithms: Nuitrack (currently used in academic research, [48–50]) and Mediapipe [51–53] (Google developed solution). The two ZED-2i cameras (2mm and 4mm lenses) were tested with the PyZED ANN algorithm [44]. All results were compared to the Vicon system, which was used as ground truth.

##### Vicon System

The Vicon system is a high quality 3D-motion capture system proven to be accurate and reliable [45,54,55]. In our experiment, 16 infrared cameras that operated at a frequency of 120 Hz simultaneously (Fig 2) were included. The optical motion capture is based on markers (infrared reflectors) that are placed by an expert on the participant’s body segments and joints [55]. Vicon Nexus software maps the camera images to a 3D coordinate system (X, Y, Z) using a direct linear transformation algorithm [56]. To identify and tag the skeleton segments and joints in the Vicon system, participants were asked to wear a body suit with 39 markers that had a radius of 14 mm, located at specific landmarks.

**Fig 2.**
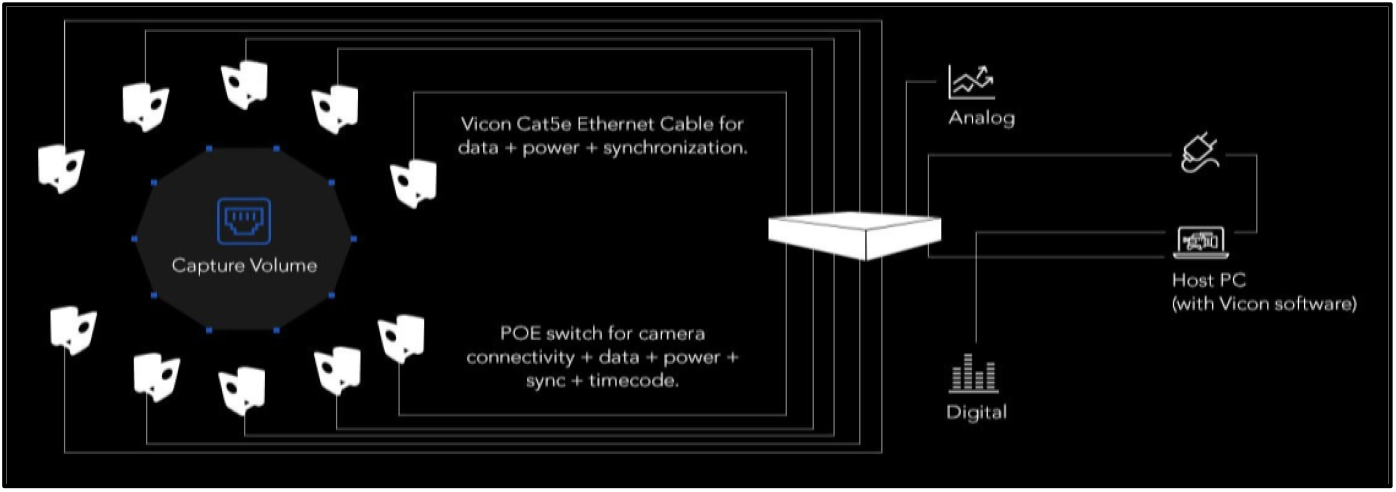
Vicon system architecture.

The obtained data were processed using the Vicon-Nexus software (version 2.5), which is considered to be the gold standard for processing, modeling, and motion analysis [5].

##### Nuitrack

Nuitrack is a software that performs skeleton tracking of users by tracking the position of 19 human joints (Fig 3) [57].

**Fig 3.**
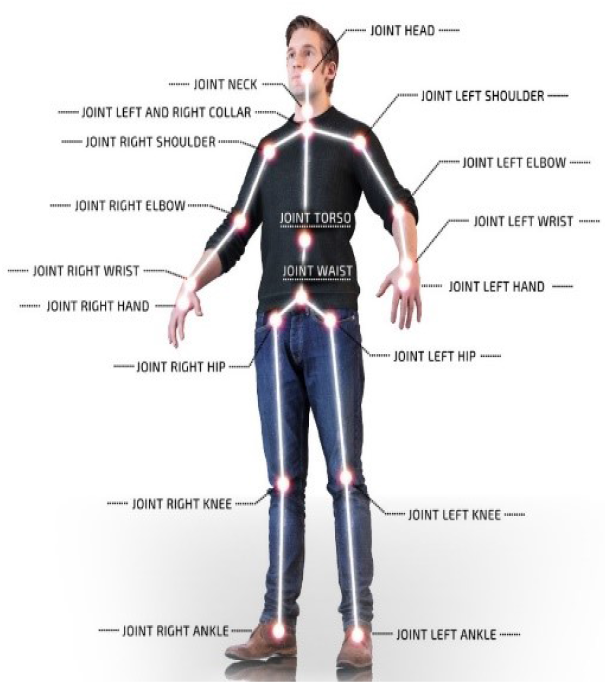
Nuitrack joints.

##### MediaPipe

MediaPipe Pose is a machine-learning (ML) solution developed by Google for high-fidelity body pose tracking, inferring 33 landmarks and a background segmentation mask on the entire body from an RGB video (Fig 4) [51].

**Fig 4.**
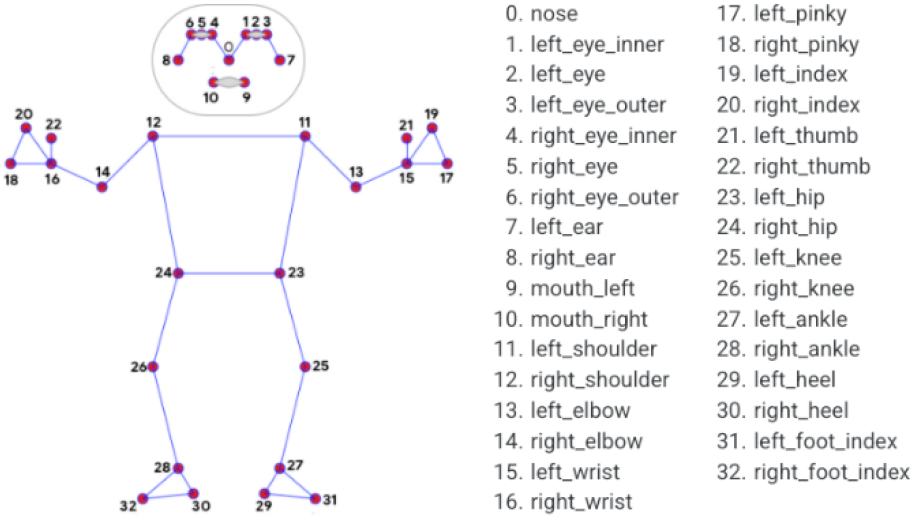
MediaPipe joints.

##### ZED-2i and PyZED

The ZED-2i is a stereo camera that provides high-definition 3D video and neural depth perception of the environment utilizing neural networks [58].

The camera is able to recognize multiple humans at the same time. It works in a dynamic state, which means the camera can move while analyzing its surroundings, and it has a working range of up to 20 meters. PyZED is a Python package that is connected to the camera and monitors 32 key points in the human body (Fig 5).

**Fig 5.**
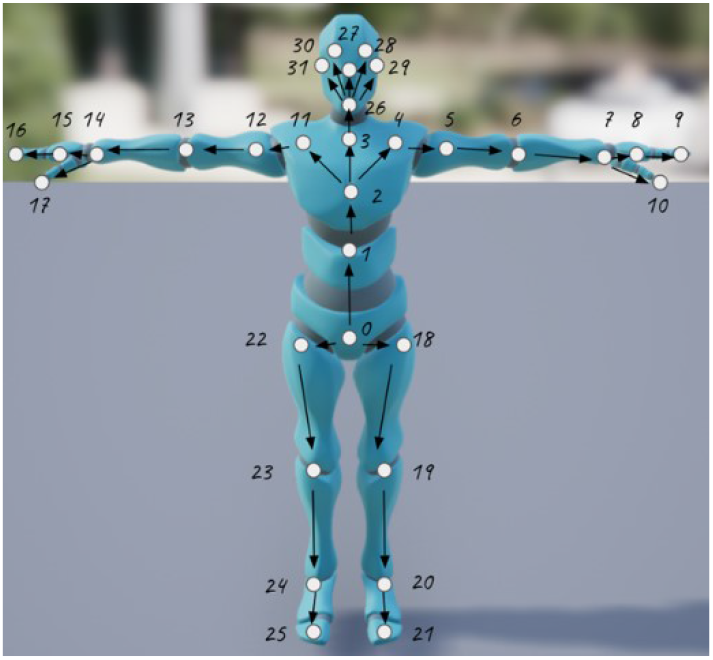
ZED-2i joints.

#### 3.1.2 Experimental Protocol

Experiments were performed in the Schwartz Movement Analysis & Rehabilitation Laboratory in the Physical Therapy Department at Ben-Gurion University of the Negev. Sixteen participants, 8 males and 8 females, 21–29 years old, participated in the experiment that included five trials. In each trial, the participant was required to walk along a 6-meter-long narrow path while moving their hands up and down horizontally. Each walk was documented by the Vicon system, a video camera, and the different cameras and algorithms. The weight, height, and arm and leg circumference of each participant were measured manually for the Vicon system (Table 2).

**Table 2.** Participant measurements.

| Parameter | Mean | Standard-deviation | Median |
| --- | --- | --- | --- |
| Height male (meters) | 1.76 | 0.064 | 1.745 |
| Height female (meters) | 1.62 | 0.058 | 1.600 |
| Weight male (Kg) | 73.930 | 8.89 | 73.0 |
| Weight female (Kg) | 53.625 | 3.77 | 53.5 |

#### 3.1.3 Experimental set-up

All four cameras (ZED2mm, ZED4mm, and two RealSense) were placed in the same position during the experiments (Fig 6). Each camera was connected to a separate computer to operate the algorithms (Table 3).

**Fig 6.**
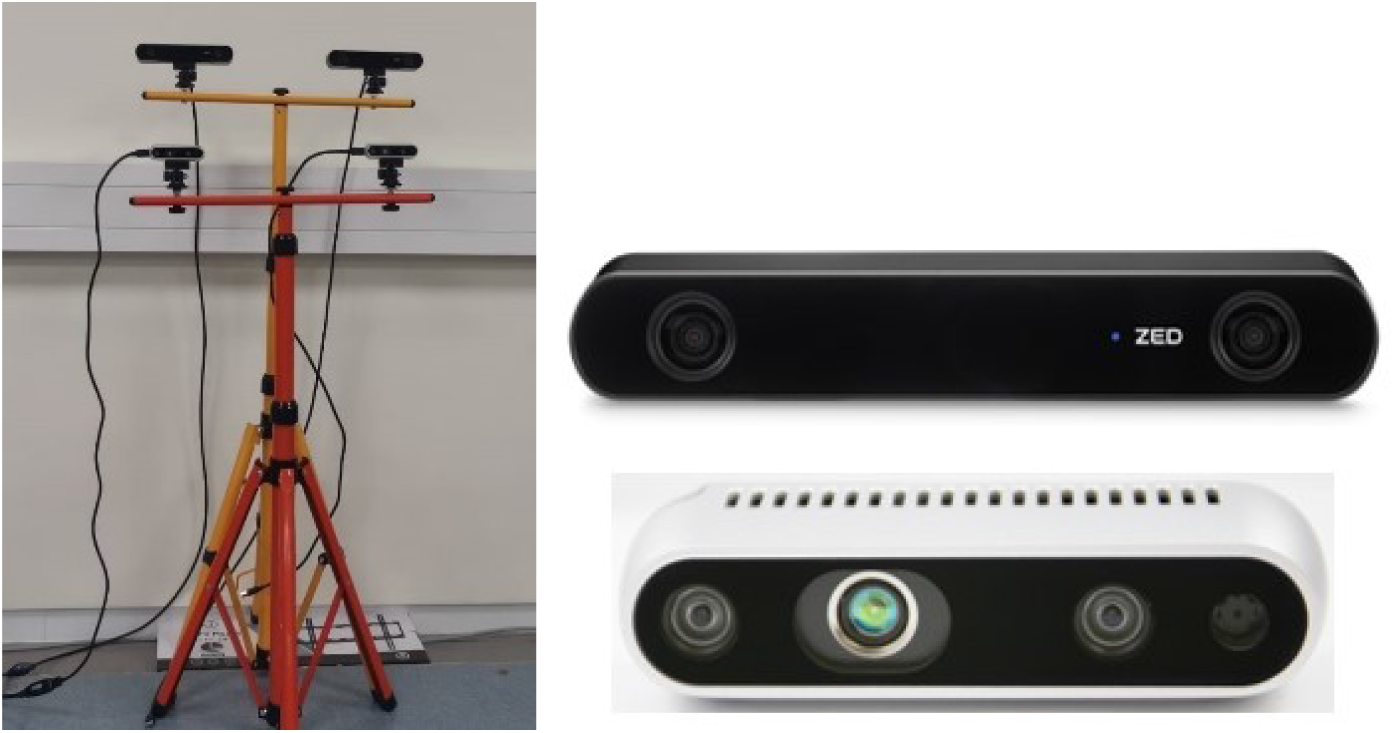
The left figure represents the camera’s position. On top are two ZED-2i cameras, on the left is a ZED2mm camera, and on the right is a ZED4mm camera. There are two RealSense cameras at the bottom. The right figure shows ZED-2i at the top and RealSense at the bottom.

**Table 3.** Technical specifications of each computer.

| Camera | Operating computer | CPU | GPU |
| --- | --- | --- | --- |
| Intel RealSense D435 | DELL Inspiron 15, 5000 Series, model: P58F001 | Intel Core i7-7500U | Intel HD Graphics 620 |
| ZED-2i | ASUS VivoBook, model:X510U | Intel Core i5-8250U | NVIDIA GeForce 930MX |
| Vicon system | DELL PC, model: Precision T3610 | Intel Xeon E5-1620 v2 | NVIDIA Quadro K600 |

During each trial, a different camera and algorithm were activated; the Vicon system was activated in each trial and served as the ground truth. In total, five trials were done per participant. One trial was carried out for each camera and algorithm (ZED2mm, ZED4mm, RS-Nuitrack, and RS-MediaPipe. In the fifth trial, two cameras (RealSense and one of the ZED-2i cameras) were used simultaneously. A total of 80 trials with all the participants were conducted (5 trials for each of the 16 participants).

#### 3.1.4 Experimental procedure

Prior to the experiment, the participants signed an informed consent form and watched a pre-filmed explanatory video of the experiment (link to the video is given in Appendix A2). After each participant’s body measurements were inserted into the system, the participant donned a special anti-bacterial suit to which the markers were attached and a headband also with markers. This was followed by a test performed to create the participant’s skeleton data for the Vicon system. Then we began the trials in random order between the four groups, as shown in Table 4. The participants were divided equally into one of the four groups. Each group included two males and two females, with a total of four participants in each group.

**Table 4.** Trial order according to algorithms.

| Group number | 1 | 2 | 3 | 4 |
| --- | --- | --- | --- | --- |
| Trial 1 | RS-MediaPipe | RS-MediaPipe +ZED2mm | ZED4mm | ZED2mm |
| Trial 2 | RS-Nuitrack | RS-MediaPipe | RS-Nuitrack +ZED4mm | ZED4mm |
| Trial 3 | ZED2mm | RS-Nuitrack | RS-MediaPipe | RS-Nuitrack +ZED2mm |
| Trial 4 | ZED4mm | ZED2mm | RS-Nuitrack | RS-MediaPipe |
| Trial 5 | RS-MediaPipe +ZED4mm | ZED4mm | ZED2mm | RS-Nuitrack |

The data were automatically recorded during each trial using the appropriate camera (RS, ZED2mm, or ZED4mm). At the end of each trial, an Excel file containing all of the segment and joint data was saved, along with the recordings of the Vicon data. This resulted in 24 files (16 trials with one camera vs. the Vicon system and eight trials with two cameras vs. the Vicon system), one for each camera and algorithm: ZED2mm, ZED4mm, RS-Nuitrack, or RS-MediaPipe from the experiment. At the end of the experiment, in order to receive information on the joint data from the Vicon system, a preliminary analysis was done using Vicon-Nexus software. Each video file was analyzed manually, with the missing joints filled in to make it as accurate as possible.

A total of 80 files were generated for the Vicon system and used to compare to each file with the other algorithms.

### 3.2 Feature extraction

#### 3.2.1 Data transformation

Each algorithm detects not only different joints but a different number of them. The following data transformations were used to compare between each joint of each specific algorithm to the body segments and joints provided by the Vicon system. Data transformations were, therefore, made at three levels.

1. Scale – the Vicon system uses mm (PyZed – meters, Nuitrack and MediaPipe – cms)
2. Rotation – current rotation in the axis to fit the Vicon system.
3. Shifting to allow comparison between the algorithms, there is a need to match their axes since each algorithm has a different absolute zero. Therefore, the Nuitrack, MediaPipe, and PyZED coordinates were shifted in accordance with Vicon coordinates. The required shifting distance was calculated for each trial separately with the most consistent joint for each of the algorithms. For the Nuitrack and PyZED, we used the CLAV joint, and for the MediaPipe, we used the mean between the right and left shoulder, which also represents the CLAV joint.

#### 3.2.2 Features

To enable the comparative analysis, the following set of features was extracted. The features were classified separately for the upper and lower extremities. Features based on the hips were calculated for both groups. Since each group of features had a different research focus, they were also evaluated separately (Table 5).

**Table 5.**
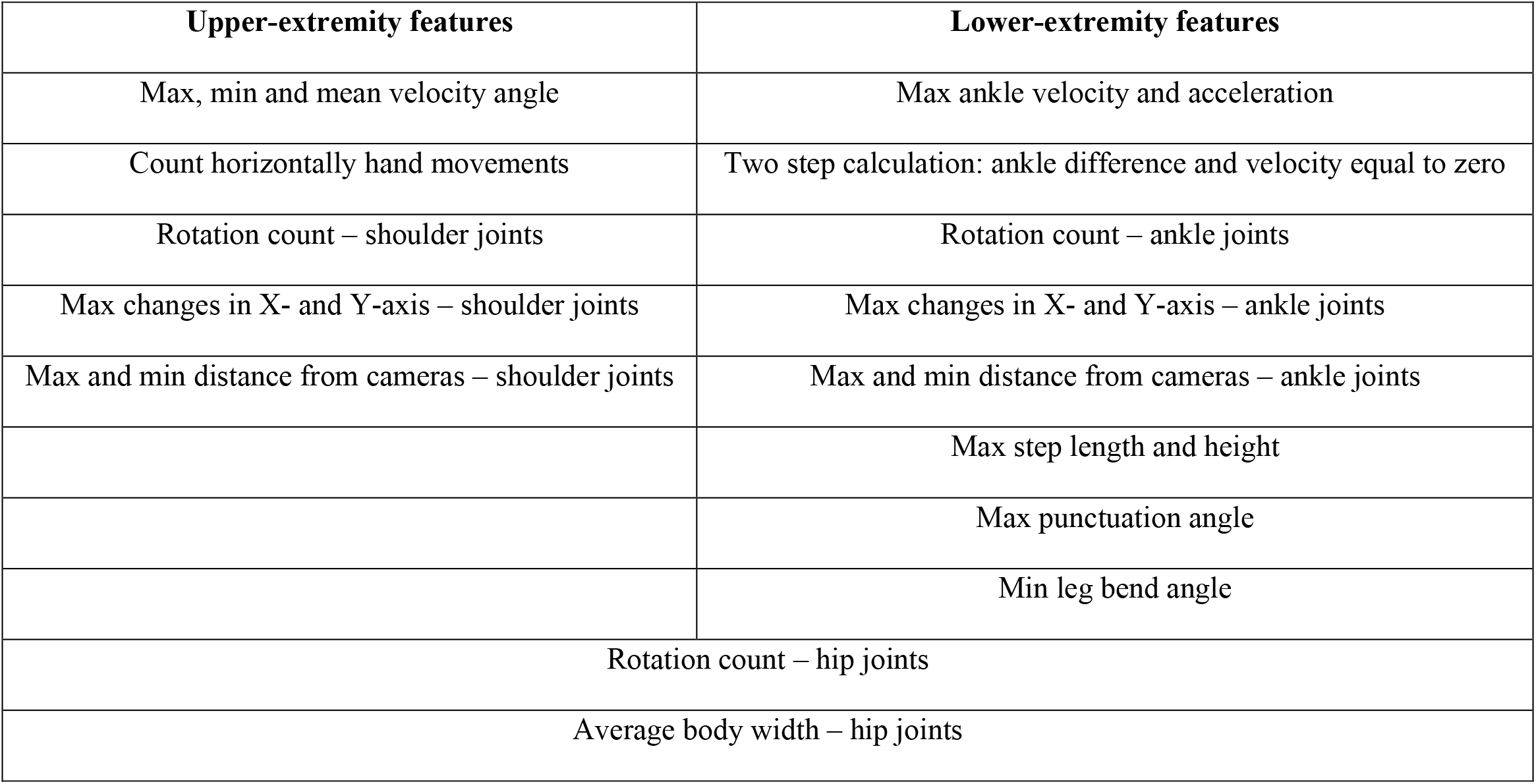
Upper- and lower-extremity features.

##### 3.2.2.1 Upper-extremity feature calculation

- Angular velocity – the deviation of the angle between the hip, shoulder, and elbow. The angle was calculated by the law of cosines:

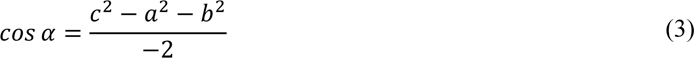

Six features were obtained from these calculations: maximum angular velocity, minimum angular velocity, and mean for each side (left and right).

- Number of horizontal hand movements – the number of repetitions based on the trend of the angle between the hands.

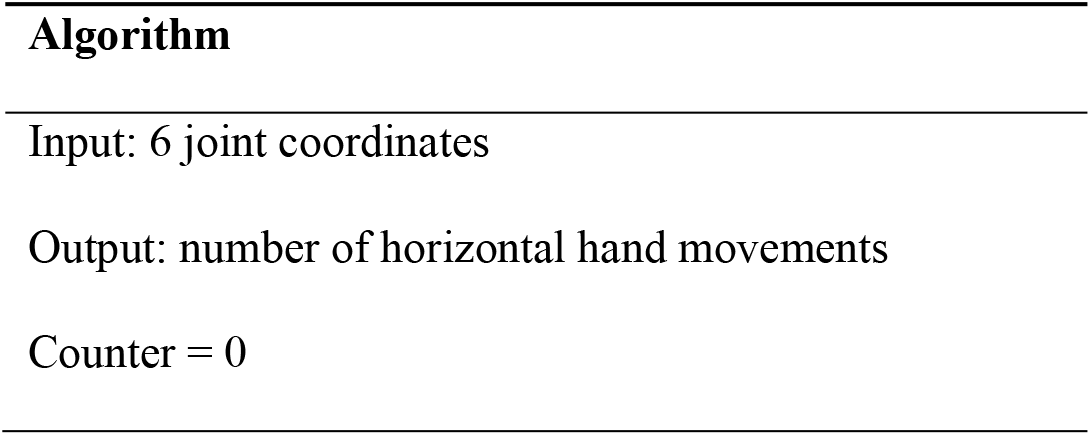
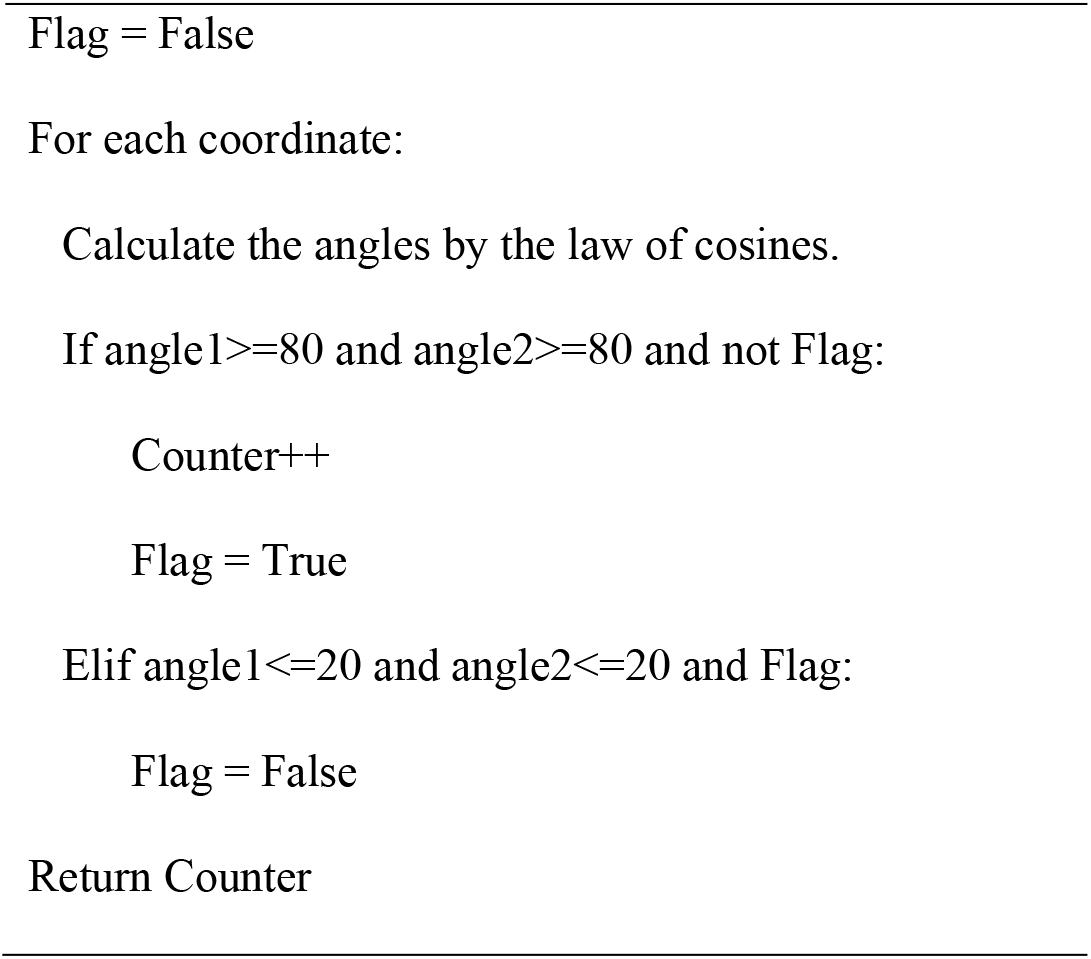
- Rotation number (the number of rotations) – the number of times the participant turned 180 degrees. This feature was calculated by comparing the X coordinates of parallel joints (hips and shoulders). When the joints’ X coordinates changed, the ratio between them was considered to be a body turn. We used this feature for two different pairs of joints: the shoulders and hips. In the experiment, each participant performed two rotations according to the given instructions.
- Average body width – For each participant, the body width was calculated according to the X and Y coordinates of the right and left hip joints, using the distance formulation between two points.

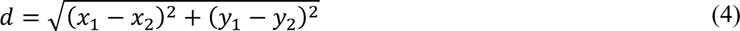

The average was calculated for all the data of body width per frame.

- Max changes in the X- and Y-axes – These features show the maximum change in shoulder position (on each side) in the X- and Y-axes. The feature is calculated by the difference between the maximum and minimum position value for a specific trial, camera and axes. It may be influenced by extreme values or camera position miscalculations.
- Max and min distance from the cameras – These features represent the maximum and minimum distance between the participant and the cameras, meaning the maximum and minimum value for the Z-axis in each trial for the shoulder joints.

##### 3.2.2.2 Lower-extremity features’ calculation

- Rotation number – the number of times the participant turned 180 degrees, as explained for the upper- extremity features. We used this feature for two different joints: the hips and ankles.
- Average body width – as explained in the upper-extremity calculations.
- Max changes in the X- and Y-axes – using the right ankle joint with the same calculation as with the upper extremity features
- Max and min distance from the cameras – using the right ankle joint with the same calculation as with the upper-extremity features
- Max ankle velocity and acceleration – The speed and acceleration were calculated using time derivations. The data were stored in four dimensions (X, Y, Z, and time) where the only axis used to calculate the velocity and acceleration was the Z-axis. Due to the discrete data sampling and the differences in acquisition rates between the cameras, the number of frames for the Vicon camera was reduced by four in order to obtain equal time differences between frames in all the cameras.
- Step count ankle difference – The number of steps is based on an algorithm that counts the number of times that the position of the ankles is met (equals). The function uses only the Z-axis for each ankle and compares the relative positions since the data are discrete. The only parameter for this algorithm was the number of frames in a row needed to decide if there was indeed a change in the position of the ankles. A grid search was done to calculate the optimal parameter value for each camera compared to the manual step count.
- Step count velocity equal to zero – The number of steps is calculated with the assumption that while taking a step, one leg stays in place and the other moves. This implies that the leg that is not in motion has zero velocity. Therefore, if we count all the times in which the derivative equals zero for each leg, we can count the number of steps taken. Since the data are continuous, there is no absolute zero value to handle this. Therefore, coarse noise filtering was added. The filter zeroed out all values that were found in the ambient delta to eliminate noise. To calculate the ambient delta, it was necessary to perform a grid search to compute the appropriate parameter for each camera.
- Max step length and height – In order to calculate these features, we used the maximum difference between the Y and Z coordinates of the right and left ankle joints.
- Max punctuation angle – In the experiment, the participants were asked to perform an exercise while remaining in a squat, changing their stance compared to the rest of the experiment (walking). This feature calculates the maximum punctuation angle for each trial and can assess the max angle calculated while the participant is in the squat trial. To calculate the punctuation angle, four joints with their X and Y coordinates are needed. These were: the right and left hip and the right and left knee.
- Min leg bend angle – This is the angle between the hip, knee, and ankle. It was calculated by the law of cosines. For this feature, we used the minimum angle.

### 3.3 Feature analysis

Every feature for each specific trial and camera with the proper algorithm was compared to the Vicon system (ground truth) by the root of the mean square error (RMSE) index (equation 1). Additionally, a linear mixed model was constructed for each individual feature to examine the coefficients and determine whether the distributions are equal. The following were inserted as the variables in equations 1 and 2 (section 2.3):

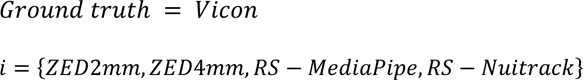

### 3.4 Ranking features

#### 3.4.1 Significant weights

The features were manually ranked in each group of features according to the relevance of the research objectives. Different rankings were compared (section 4.1.1). The lowest rank (1) in each group of features is assigned to the least significant feature, while the highest rank is assigned to the most significant feature. We calculated the weight of each feature by dividing its rank by the sum of its ranking in each group. In Table 6, the features are ranked according to their significant in each of the groups, and weighted by significant, where the sum of all weights multiplied by the features amount is 100%. The specific rankings were selected by the researchers (and can be changed).

**Table 6.** The features and the significant weights. The amount column represents the number of features in a specific row.

| Upper extremity |  |  |  | Lower extremity |  |  |  |
| --- | --- | --- | --- | --- | --- | --- | --- |
| Features | Rank | Amount | Weight% | Features | Rank | Amount | Weight% |
| Max, min and mean velocity angle | 6 | 6 | 8.96 | Max ankle velocity and acceleration | 4 | 2 | 7.54 |
| Count horizontal hand movements | 7 | 1 | 10.45 | Two-step calculation: ankle joint difference and velocity equal to zero | 6 | 2 | 11.33 |
| Rotation count – shoulder joints | 5 | 1 | 5.97 | Rotation count – ankle joints | 1 | 1 | 1.90 |
| Rotation count – hip joints | 4 | 1 | 7.46 | Rotation count – hip joints | 1 | 1 | 1.90 |
| Average body width – hip joints | 3 | 1 | 4.48 | Average body width – hip joints | 1 | 1 | 1.90 |
| Max changes in X- and Y-axis – shoulder joints | 2 | 4 | 2.99 | Max change in X-axis – ankle joint | 4 | 1 | 7.54 |
| Max and min distance from cameras – shoulder joints | 1 | 4 | 1.48 | Max change in Y-axis – ankle joint | 2 | 1 | 3.76 |
|  |  |  |  | Max and min distance from cameras – ankle joints | 6 | 2 | 11.33 |
|  |  |  |  | Max step length and height | 4 | 2 | 7.54 |
|  |  |  |  | Max punctuation angle | 2 | 1 | 3.76 |
|  |  |  |  | Min leg bend angle | 2 | 1 | 3.76 |

#### 3.4.2 Grading techniques

The grading techniques are used to create the same scale for the RMSE index in all features so they will be comparable. In this demonstration, two different grading techniques were used to compare the camera performances for each feature using the RMSE index, namely basic normalization and relative normalization.

In the first technique, basic normalization: the camera that gained the lowest RMSE value per feature was graded as 0, and the camera that gained the highest RMSE value was graded as 3. With this technique, we create an ordinal scale avoiding the impact of the RMSE on different values per feature.

In the second technique, relative normalization, the RMSE values were transformed to the segment [0,1] using equation 5.

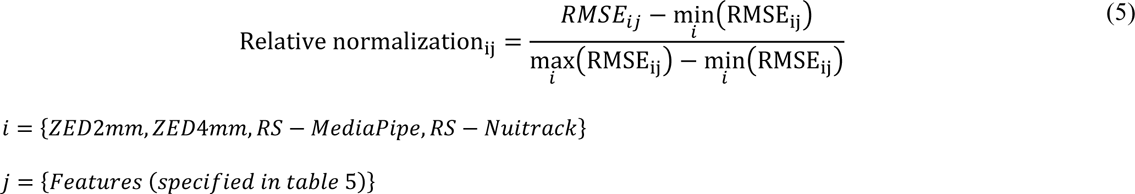

Relative normalization is a type of calibration that considers the effect of the size of each RMSE value.

For each grading technique, the RMSE grades should be summed per camera and algorithm.

## 4 Results

### 4.1 RMSE

The results of each trial with each camera were compared to the Vicon ground truth to calculate RMSE index values. The first step was to analyze the results for equal weights without significant features. Comparing the RMSEs with the two grading techniques showed that the ZED cameras had lower RMSE values than the RS- Nuitrack and RS-MediaPipe in most cases (about 60% for the upper and lower extremity features).

For the upper extremity, the average value of ZED2mm was 158.42% better than the next performing camera (ZED4mm) and 338.29% better as compared to the poorest performing camera (RS-MediaPipe) based on the average percentage between the basic and relative normalizations. For the lower extremity, the average value of the ZED2mm was better by 183.845% than the next performing camera (ZED4mm) and by 367.135% as compared to the worst performing camera (RS-MediaPipe) for the average percentage between the basic and relative normalizations (Table 7).

**Table 7.** The average, standard deviation, and median for each camera’s RMSE index with the two grading techniques.

| Upper extremity |  |  |  |  |  |  |  |  |
| --- | --- | --- | --- | --- | --- | --- | --- | --- |
|  | Basic normalization |  |  |  | Relative normalization |  |  |  |
|  | ZED2mm | ZED4mm | RS-MediaPipe | RS-Nuitrack | ZED2mm | ZED4mm | RS-MediaPipe | RS-Nuitrack |
| Average | 0.888 | 1.055 | 2.166 | 1.888 | 0.153 | 0.303 | 0.662 | 0.556 |
| Standard deviation | 0.936 | 1.025 | 0.763 | 1.149 | 0.256 | 0.378 | 0.352 | 0.427 |
| Median | 1 | 1 | 2 | 2 | 0.018 | 0.137 | 0.668 | 0.551 |
| Lower extremity |  |  |  |  |  |  |  |  |
|  | Basic normalization |  |  |  | Relative normalization |  |  |  |
|  | ZED2mm | ZED4mm | RS-MediaPipe | RS-Nuitrack | ZED2mm | ZED4mm | RS-MediaPipe | RS-Nuitrack |
| Average | 0.8 | 1.2 | 2.266 | 1.733 | 0.147 | 0.32 | 0.663 | 0.518 |
| Standard deviation | 0.832 | 1.045 | 0.928 | 1.062 | 0.259 | 0.364 | 0.401 | 0.400 |
| Median | 1 | 1 | 3 | 2 | 0.056 | 0.163 | 1 | 0.464 |

To assess the cameras’ performances with the proper algorithm across all features, we sum the RMSE results according to the basic and relative normalization (Fig 7). The best overall performance is represented by minimum value. The results in Fig 7 in the case of equal weights for each feature reveal that the ZED2mm resulted in the best score and the RS-MediaPipe resulted in the worst result for both basic and relative normalizations for both groups of features.

**Fig 7.**
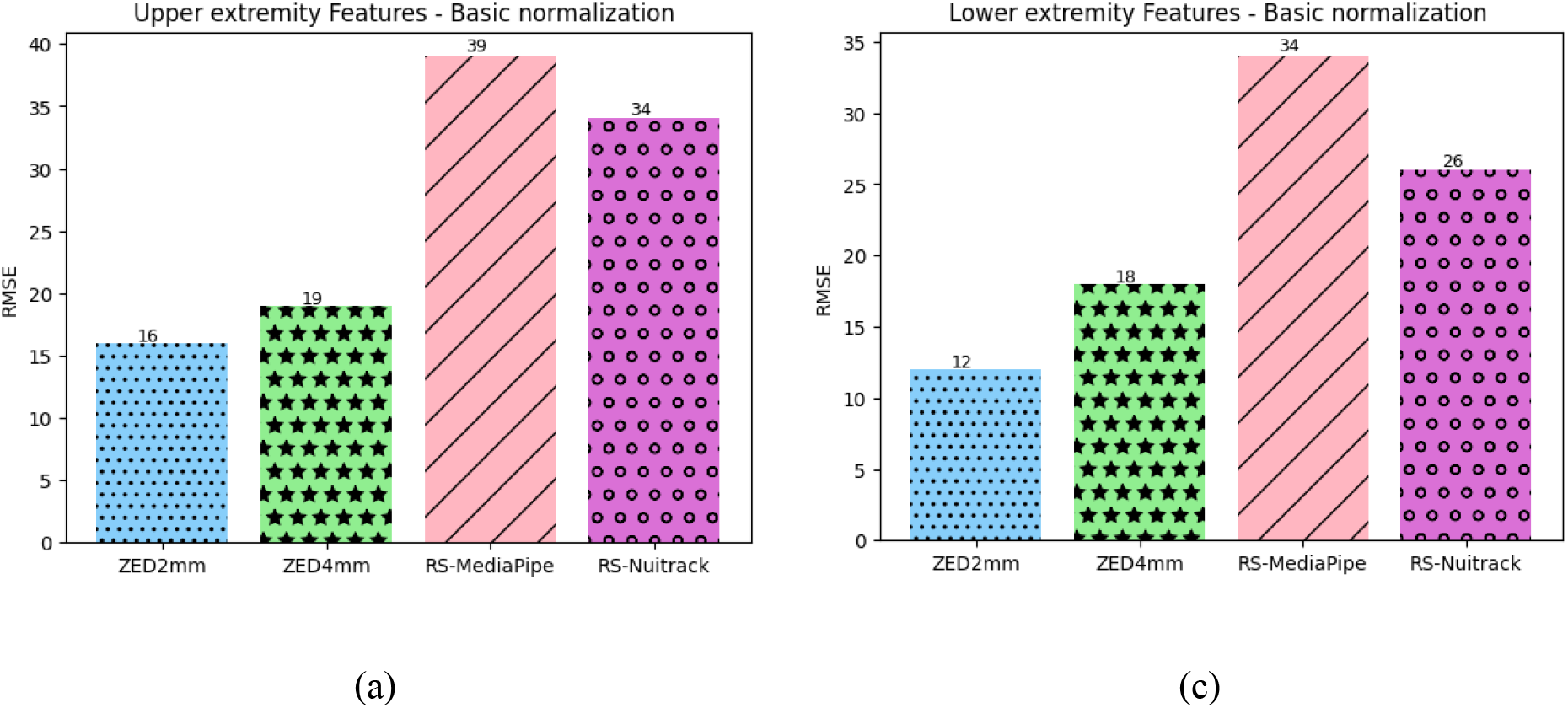
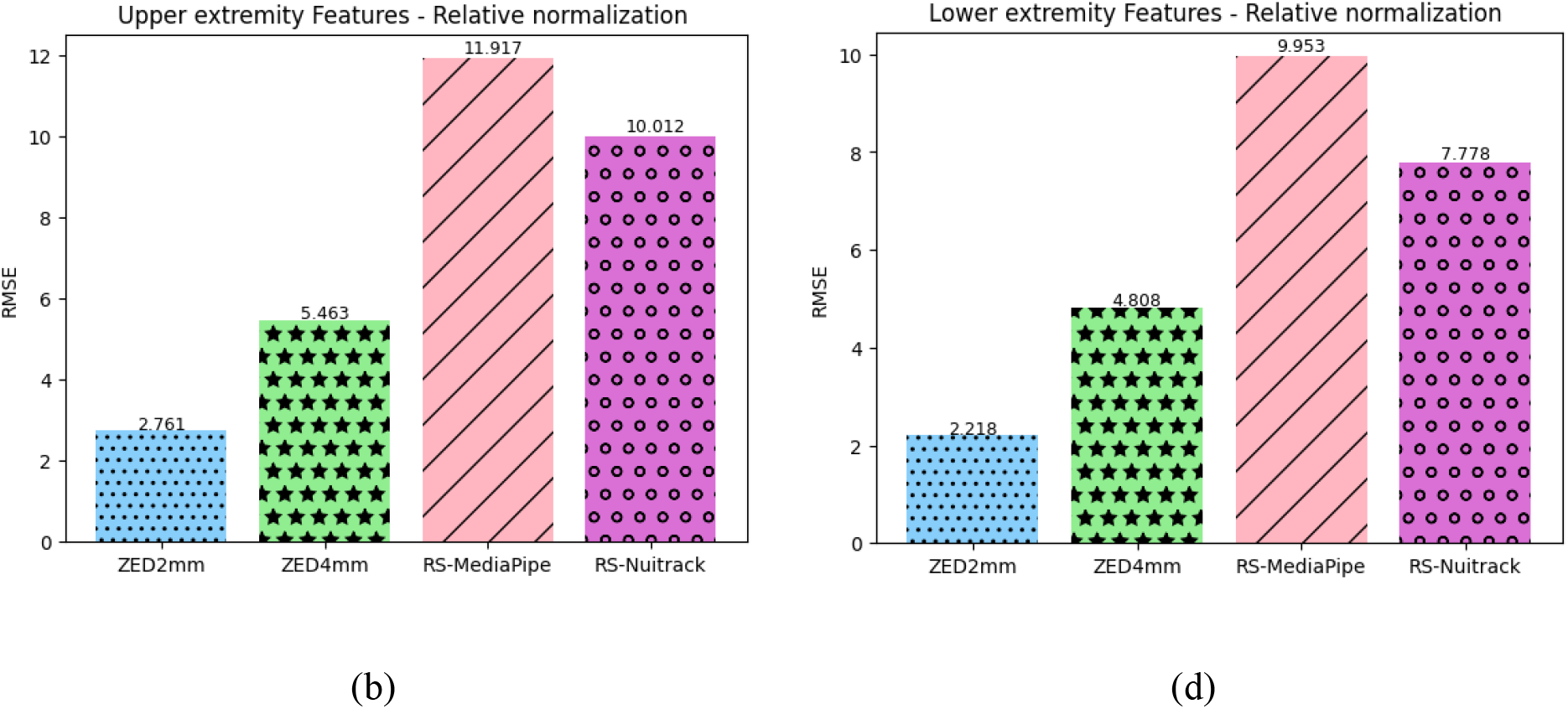
The sum of RMSE results for equal weights for all features: (a) basic and (b) relative normalization for upper-extremity results; (c) basic and (d) relative normalization for lower-extremity results.

The second step was to analyze the result for different significant weights, as described in Table 6. The results (Fig 8) reveal different results for the two groups of features for different significant weights of features. For the lower extremity, the ZED2mm resulted in the best score of 572.8% compared to the RS-MediaPipe, which had the worst result in the average between basic and relative normalizations. For the upper extremity, the best camera difference was between basic and relative normalizations (Table 8). For the basic normalization, the camera with the best result was the ZED4mm by 232.68% compared to the RS-Nuitrack, which had the worst result. For the relative normalization, the camera with the best result was the ZED2mm by 318.88% compared to the RS-Nuitrack, which had the worst result.

**Fig 8.**
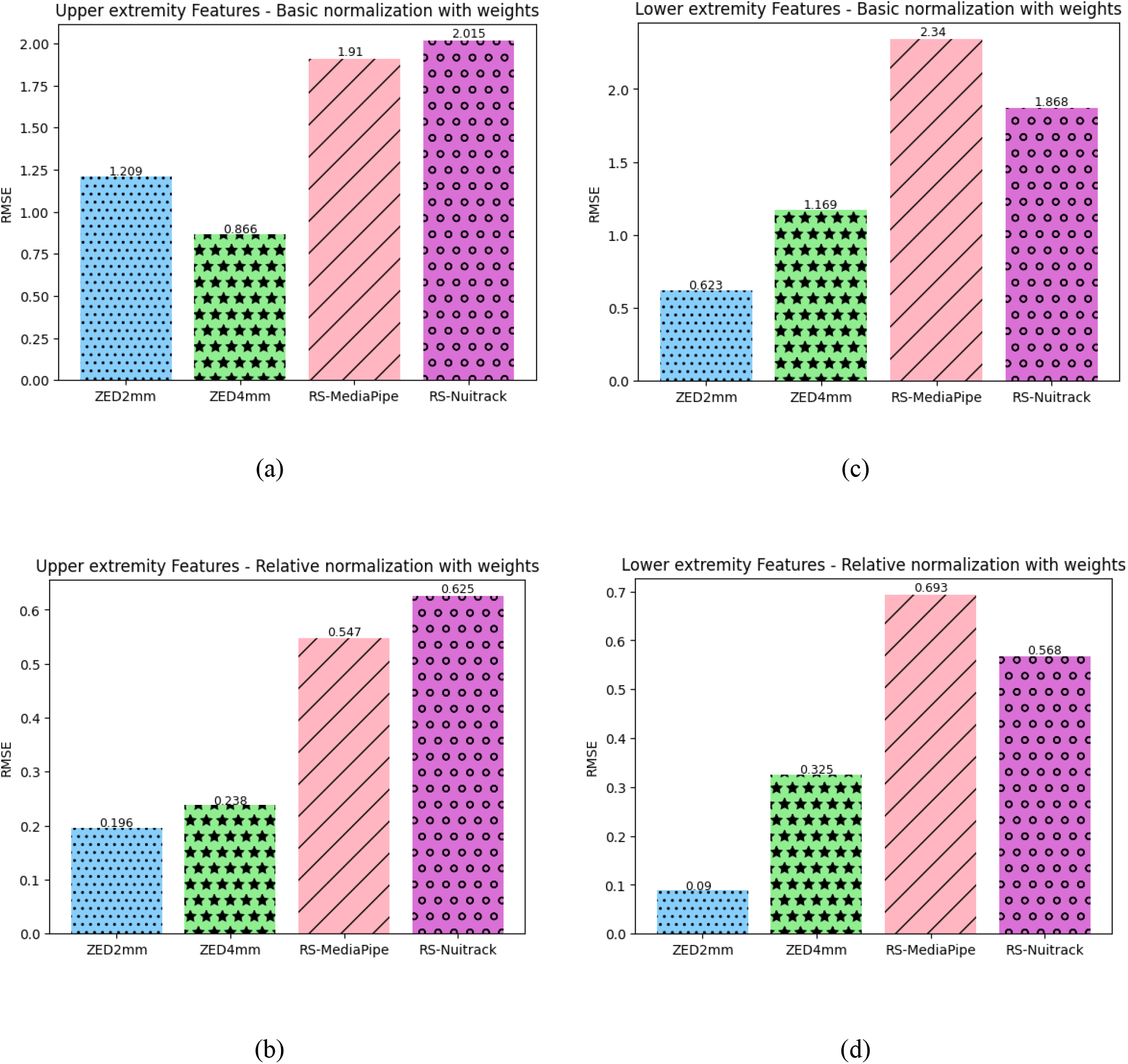
The sum of RMSE results with the significance of feature weights: (a) basic and (b) relative normalization for upper-extremity results; (c) basic and (d) relative normalization for lower-extremity results.

**Table 8.** The results for each group of features with the significance of feature weights.

| Upper extremity | Lower extremity |
| --- | --- |
| ZED4mm (basic normalization), ZED2mm(relative normalization) | ZED2mm |
| ZED4mm (relative normalization), ZED2mm(basic normalization) | ZED4mm |
| RS-MediaPipe | RS-Nuitrack |
| RS-Nuitrack | RS-MediaPipe |

The RS-MediaPipe and RS-Nuitrack produced the poorest results for both the upper- and lower-extremity features, probably due to their lack of depth perception. The RS-MediaPipe camera uses RGB video frames to identify people and their location in space, where each frame has only one RGB image [51], whereas the ZED-2i takes two RGB images per frame, giving it more data to evaluate depth in space better. The Nuitrack uses the RealSense laser (infrared) sensor and the RGB camera to evaluate depth in space. While laser sensing is good for identifying an object’s location, it is sensitive to glare, as once the laser beam hits a certain part, it reflects light in all directions. This implies that the detection of nearby or moving objects is not accurate for this technology. For example, the laser reflection from the left leg will affect the reflection from the right leg when standing toward the camera.

The ZED4mm is intended for image processing from a large distance, and the ZED2mm is suitable for close-to-medium distances. The specified distances for each lens are not specified in the camera descriptions and are denoted as 0.2–20 meters for both. The ZED4mm has a smaller field of view (72°H x 44°V x 81°D max) compared to the ZED2mm (110°H x 70°V x 120°D max), implying that when the Z-axis is used, the ZED4mm performance was not as good as that of the ZED2mm for this specific experiment and the features selected. The feature’s importance in the upper and lower extremities affected the results. In the lower extremity, the most important features refer to the Z-axis and, therefore, to its higher ranking. In the upper extremities, there was a difference between the basic and relative normalization in the first and second places. This difference is due to the fact that the ZED4mm has more features with a lower RMSE index than the ZED2mm, yielding a better performance for the ZED4mm. While the ZED2mm had the lowest RMSE, in some cases, the ZED4mm had a high RMSE, which affected the relative normalization significantly.

#### 4.1.1 Sensitivity analysis

A sensitivity analysis for different weights revealed that results depend on the significant of the features, revealing the importance of this analysis. Different weights may yield varying scores for each camera-algorithm couple.

However, in our demonstration for the different weights analyzed, the best and second-best performance was between the ZED cameras, and the third best and worst performance was between the Nuitrack and MediaPipe cameras (Table 9).

**Table 9.**
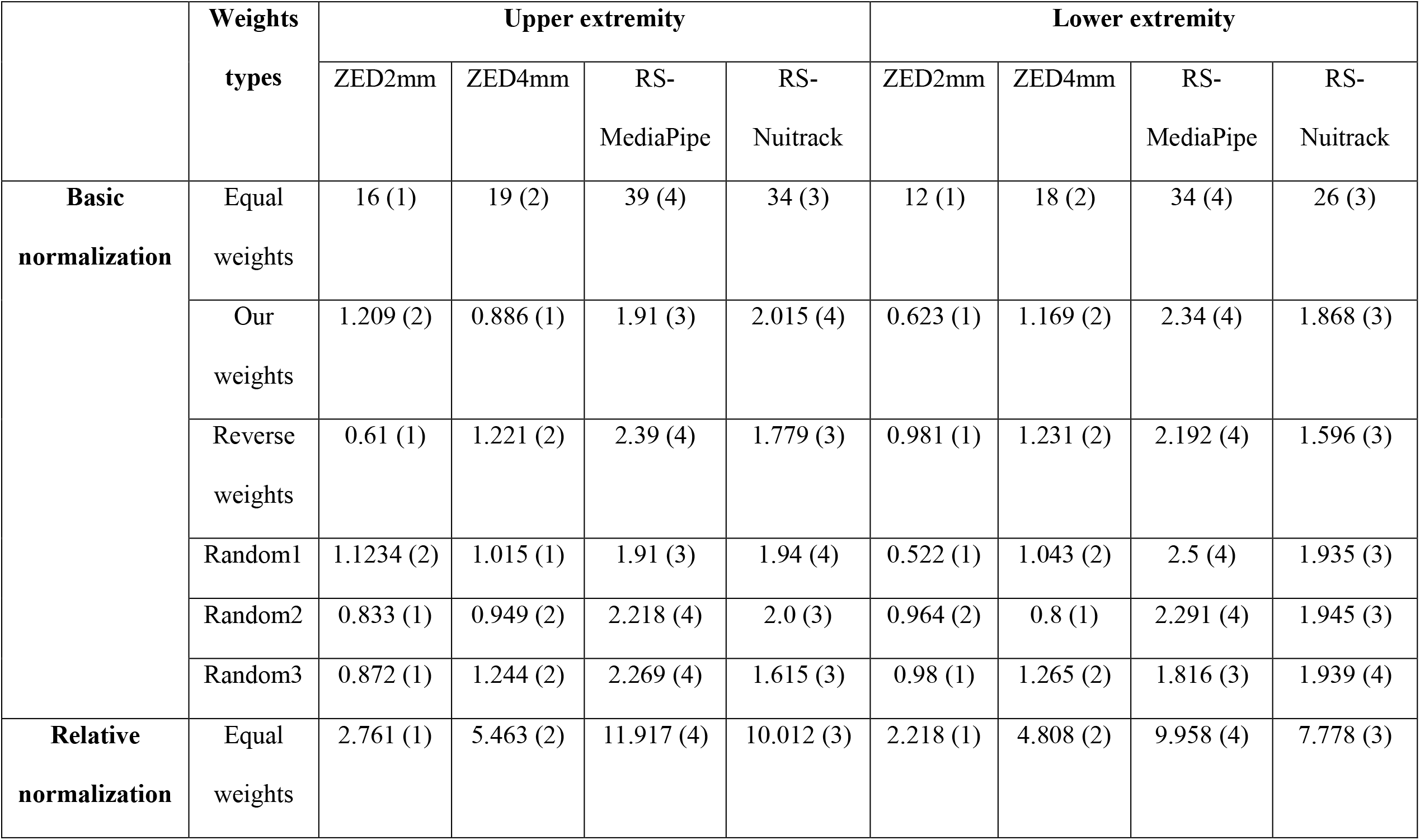
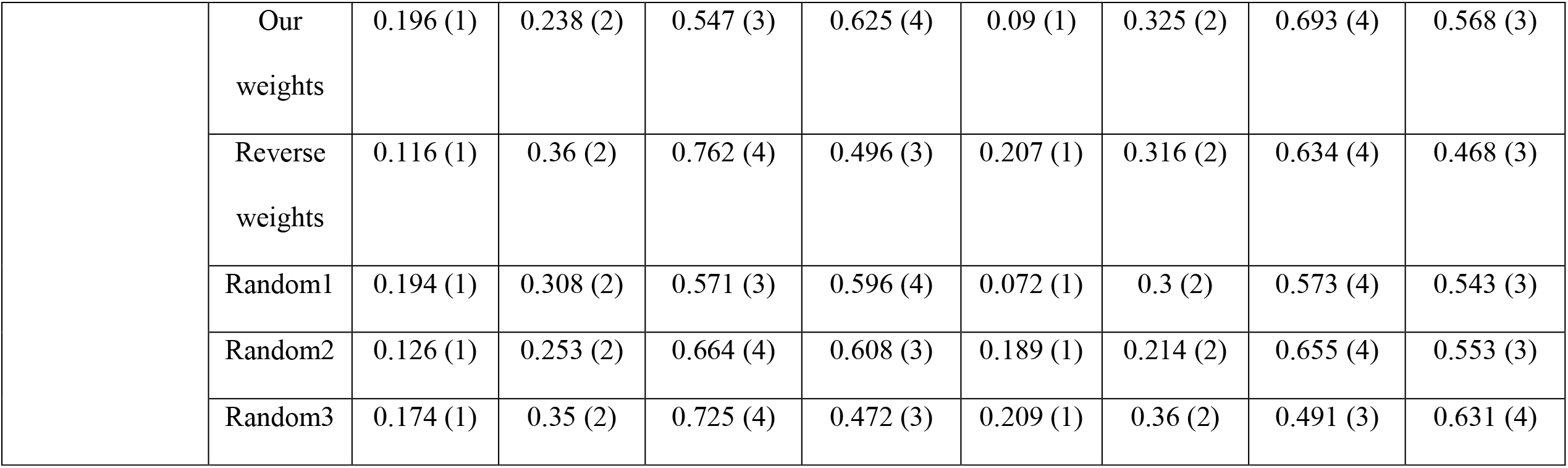
Sensitivity analysis results for each group. The value listed in the table is the RMSE index, and in brackets, the rank according to all the cameras.

We also performed a sensitivity analysis for all of the features combined (Table 10, in case of duplication, duplicates were removed in some features).

**Table 10.** Sensitivity analysis results for all features combined.

| Both groups |  |  |  |  |  |
| --- | --- | --- | --- | --- | --- |
|  | Weights types | ZED2mm | ZED4mm | RS-MediaPipe | RS-Nuitrack |
| Basic normalization | Equal weights | 24 (1) | 35 (2) | 69 (4) | 58 (3) |
|  | Random1 | 0.703 (1) | 1.197 (2) | 2.376 (4) | 1.725 (3) |
|  | Random2 | 0.952 (2) | 0.757 (1) | 2.217 (4) | 2.074 (3) |
|  | Random3 | 0.797 (1) | 0.929 (2) | 1.962 (3) | 2.313 (4) |
| Relative normalization | Equal weights | 3.646 (1) | 9.414 (2) | 20.315 (4) | 17.326 (3) |
|  | Random1 | 0.099 (1) | 0.304 (2) | 0.695 (4) | 0.489 (3) |
|  | Random2 | 0.143 (1) | 0.172 (2) | 0.651 (4) | 0.61 (3) |
|  | Random3 | 0.087 (1) | 0.241 (2) | 0.513 (3) | 0.731 (4) |

Sensitivity analyses (Table 11) summarize the main findings. For the basic normalization, the ZED2mm outperformed the ZED4mm by three times (the best performance of all), while for relative normalization, the ZED2mm always performed better than the ZED4mm. Compared to MediaPipe, Nuitrack performed three times better in basic and relative normalization.

**Table 11.** Summary of sensitivity analyses results.

|  |  | 1 | 2 | 3 | 4 |
| --- | --- | --- | --- | --- | --- |
| Basic normalization | ZED2mm | 12 | 4 | - | - |
|  | ZED4mm | 4 | 12 | - | - |
|  | RS-MediaPipe | - | - | 4 | 12 |
|  | RS-NuiTrack | - | - | 12 | 4 |
| Relative normalization | ZED2mm | 16 | - | - | - |
|  | ZED4mm | - | 16 | - | - |
|  | RS-MediaPipe | - | - | 4 | 12 |
|  | RS-NuiTrack | - | - | 12 | 4 |

### 4.2 Linear mix model results

In the first stage, the model construction method was based on the feature type, which was divided into two groups: normal distribution and discrete (count horizontal hand movements, step calculation, and rotation count). For the features with normal distribution, we performed a regular linear mixed model, and for the features with discrete distribution, we performed a generalized linear mixed model with the Poisson family and log link function.

The outcome of the models was the 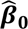, which is the estimated intercept, and 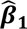, which is the estimated regression coefficient (slope) for the independent variable (feature values of the examined depth camera and algorithm). In the second stage, we performed a Wald test with the null hypothesis that 𝜷_𝟏_, the regression coefficient, is equal to 1. Accepting the null hypothesis implies that the examined feature by the camera and algorithms had the same trend when compared to the Vicon results. Table 12 represents the features that had a significant similarity to the Vicon results for the upper and lower extremity features and are therefore deemed to be good indicators of the measured outcome.

**Table 12.**
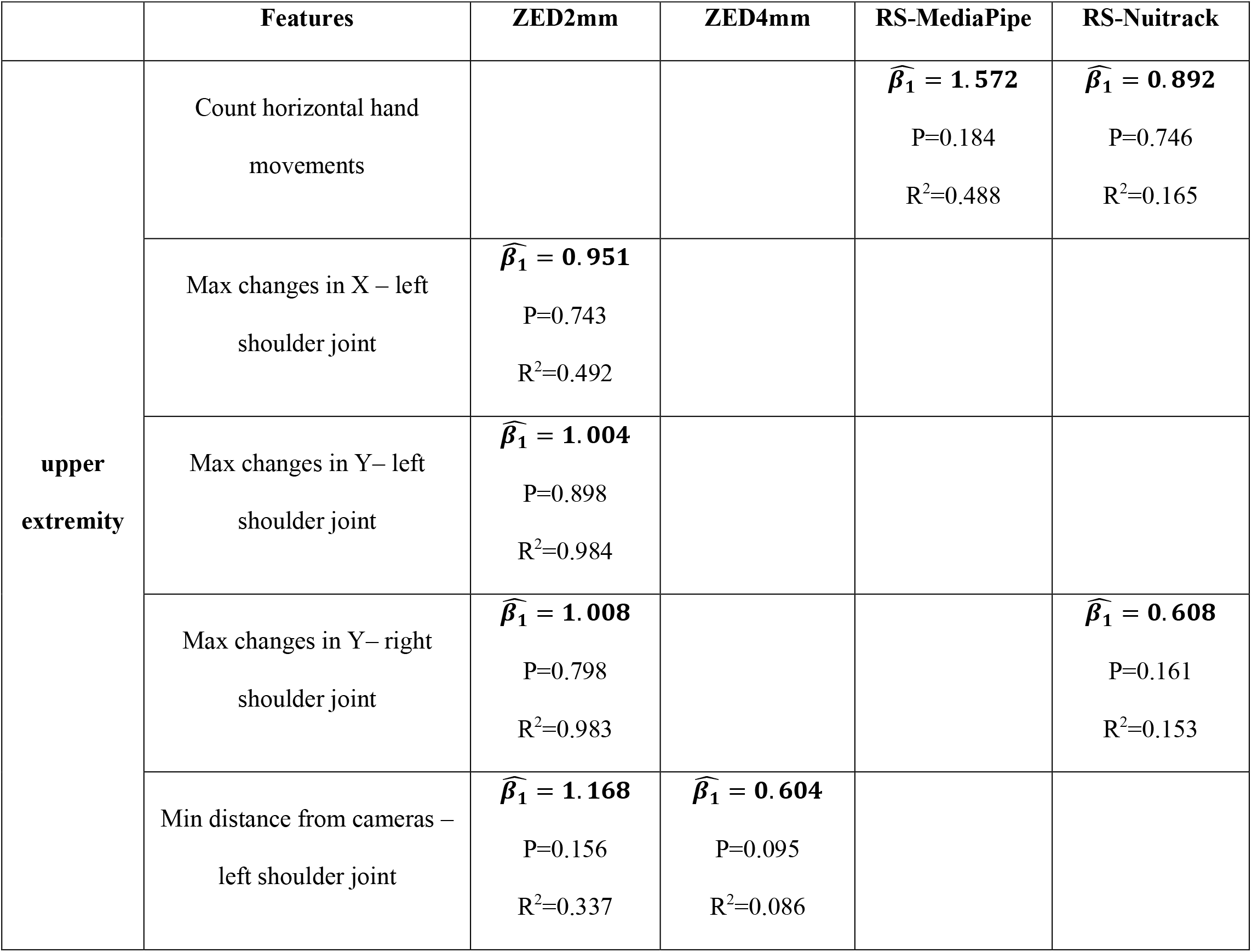
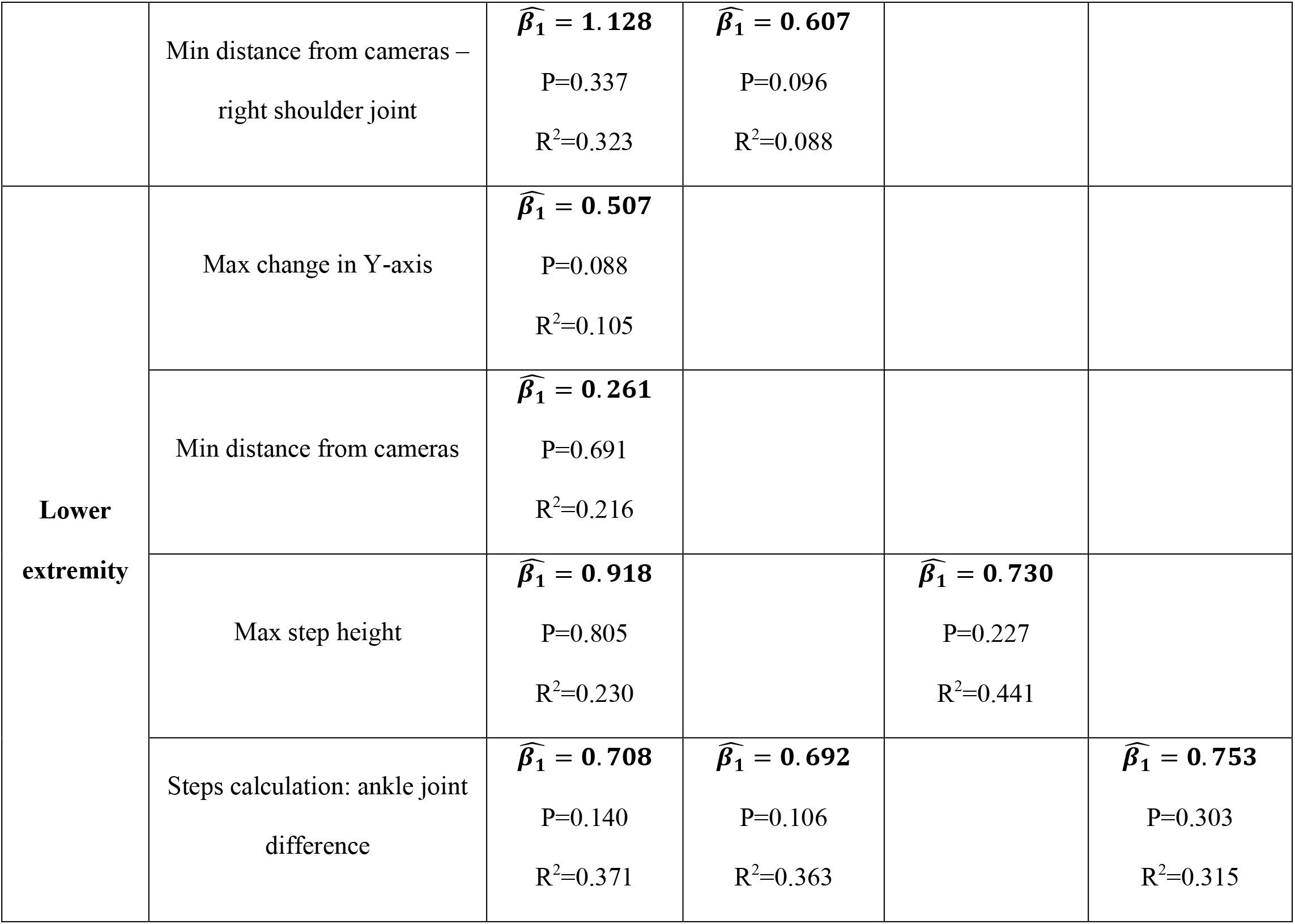
The result of the Wald test. P represents the P-value of the test that the regression coefficient, is equal to 1. R^2^ represents the marginal R^2^ of the model, the goodness-of-fit (model accuracy), without taking the random effect in the prediction.

The results (Table 12) show that the ZED2mm camera performed better than the other cameras and algorithms for all the features that were not rejected in the null hypothesis, with six features in the upper extremity and four features in the lower extremity.

For the upper extremity, the results show that the RealSense camera with either the Nuitrack or MediaPipe algorithm was the preferred choice for the most important feature (count hands movement). However, when we look at the R^2^ in the features “max changes in Y” with both sides, the ZED2mm performed the best, and the model accuracy is very high.

For the lower extremity features, the ZED2mm camera performed better, particularly for the features with high importance, and the min distance from the cameras and step counts with ankle difference features have high weight and importance.

## 5 Discussion and conclusion

Our research presents a comparative analysis methodology with four main steps for evaluating cameras and software tools utilized in skeleton tracking as well as comparing these methodologies to ground truth. The methodology describes the experimental design and analysis methods for comparing both upper and lower extremities.

The methodology was demonstrated on three cameras and three algorithms. The findings suggest that the ZED2mm camera would be a suitable choice for skeleton tracking and analyzing human activity across the entire body, based on both analysis of the RMSE results and linear mixed models. For both upper- and lower-extremity features, the ZED-2i cameras produced better performance when compared to the RealSense camera with the Nuitrack and MediaPipe software according to the RMSE index.

A sensitivity analysis for different weights indicates that the ZED2mm had the best score for the majority of the weight’s types for both upper- and lower-extremity features and their combination. This indicates that the ZED-2i cameras provide comparable data to a ground truth 3D motion analysis system when assessing upper- and lower-extremity features, as well as balance and gait. This could provide geriatric and rehabilitation practitioners with important information that is not currently obtainable in the clinical setting.

However, the choice of camera and algorithm ultimately depends on the specific requirements of the application and the environment in which the analysis will take place. Hence, varying weights and preferences in each research study emphasize the need for customized camera selection and algorithm depending on the research objective. This ensures that the desired/accurate outcomes and meaningful insights are achieved effectively.

The comparative analysis methodology provides a systematic way to compare software tools and cameras for skeleton tracking. The ranking method enables to prioritize different features depending on the research objective.

## Acknowledgments

This project was partially supported by Ben-Gurion University of the Negev through the Agricultural, Biological and Cognitive Robotics Initiative, the Marcus Endowment Fund, and the W. Gunther Plaut Chair in Manufacturing Engineering.

The authors thank the volunteers who participated in this study.

## Institutional Review Board Statement

“The study was conducted in accordance with the Declaration of Helsinki and approved by the Ethics Committee of the Ben-Gurion University of the Negev”.

## Appendix A

### A1. Code

Code Share - Comparative analysis. Available :https://github.com/AMeshu25/Comparative_analysis.git

### A2. Video

Video of experiment description.

Available: https://drive.google.com/file/d/10p6Ngsade3IOS-Wigcq4kAGW9wJ1RPYR/view?usp=drive_link

